# Multi-objective optimization for designing structurally similar proteins with diverse sequences

**DOI:** 10.1101/2025.09.13.676063

**Authors:** Ryo Akiba, Yoshitaka Moriwaki, Ryuichiro Ishitani, Naruki Yoshikawa

## Abstract

Recent advances in artificial intelligence technologies have accelerated the development of computational protein design techniques. Although the structure of the designed proteins has been the primary focus, diversity of designed amino acid sequences is another important aspect of protein design. To address the trade-off between reducing sequence similarity and improving structural similarity, simultaneous optimization of these two objectives can be effective. We present a method that integrates ProteinMPNN with the multi-objective optimization algorithm NSGA-II to design proteins that retain high structural similarity to a target while exhibiting low sequence similarity. Using Top7 as a reference protein, we demonstrate that our approach improves sequence diversity compared to the original ProteinMPNN while maintaining comparable structural similarity.

## 1. Introduction

Protein design is an important research area that can be applied to diverse applications, such as drug discovery and enzyme development. With recent advances in artificial intelligence (AI), various generative AI methods for protein design have been proposed [1]. Among these, two-staged protein design methods have demonstrated superior performance and attracted considerable attention. They divide the protein design process into the generation of three-dimensional structure and the generation of amino acid sequences that fold into the designed structure.

In the first stage of structural design, diffusion models, a type of deep generative model, are widely used [2]. Diffusion-based methods such as RFDiffusion [3] and Chroma [4] generate diverse protein backbones by gradually transforming random noise into meaningful structures. Another approach, known as the hallucination method [5,6], searches for structures that maximize an objective function using structure prediction models.

In the second stage of sequence design, conventional approaches have relied on physics-based tools that search for sequences based on energy calculations [1]. More recently, machine learning methods using graph neural networks have emerged. For example, ProteinMPNN (pMPNN) [7] employs a message passing neural network that represents protein backbones as graphs, where interatomic distances and orientations are encoded as features. This enables the model to predict amino acid sequences that fold into structures similar to a given backbone. Compared with traditional physics-based methods, pMPNN achieves higher sequence recovery for protein backbones. Moreover, its designs have been experimentally validated, confirming that the generated sequences robustly produce the target structures [3,6].

However, pMPNN tends to generate sequences that cluster within narrow regions of sequence space. This lack of diversity limits exploration of the broader landscape and hinders the discovery of proteins optimized not only for structure but also for additional properties, such as stability and solubility, which are critical in practical applications. To improve the diversity of designed proteins, several machine learning methods have been proposed [8–10].

Since proteins with similar amino acid sequences tend to have similar structures, there is a trade-off between reducing sequence similarity and improving structural similarity. Multi-objective optimization [11] is often employed to simultaneously optimize multiple objective functions in trade-offs. Recently, several studies have explored the introduction of multi-objective optimization algorithms into protein sequence design. For example, Luo et al. proposed a deep learning-guided algorithm that designs amino acid sequences that jointly optimize multiple experimental properties, such as fluorescence intensity and stability, or binding affinities to multiple targets [12]. Hong et al. integrated AI models into the NSGA-II framework and successfully optimized the stability of switch proteins in different conformational states, as well as the stability of proteins in various binding states [13]. However, these studies primarily focus on improving protein performance, and sequence diversity was not explicitly considered.

In this study, we propose a protein structure design method that combines pMPNN with the multi-objective optimization algorithm NSGA-II [14] to simultaneously optimize two objectives in trade-off: sequence dissimilarity and structural similarity. Using the TOP7 protein as a test case, our method successfully designed novel proteins that are structurally similar to the original protein while exhibiting low sequence similarity, with less than 10% of amino acids identical to the original sequence. These results demonstrate that the performance of generative AI models for protein design can be improved by combining them with multi-objective optimization methods.

## 2. Method

In this study, we propose an algorithm that combines the protein design method pMPNN, which generates sequences that fold into a given structure, with the multi-objective optimization algorithm NSGA-II to enable the design of proteins that simultaneously improve multiple evaluation criteria.

### 2.1. Multi-objective optimization

The problem of simultaneously optimizing multiple objective functions is called a multi-objective optimization problem [11], and it can be formulated as Equation (1).

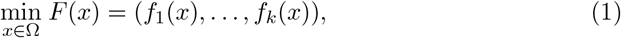

where Ω is the domain of the variable *x, k* ≥ 2 is the number of objective functions, and *f*_*i*_(*x*) (*i* = 1, …, *k*) are objective functions.

Since there is no single solution that simultaneously optimizes all conflicting objective functions, the goal of multi-objective optimization is to find a set of solutions that are not dominated by any other feasible solutions. The variable *x* is said to dominate another variable *x*^*′*^ when Equation (2) holds.

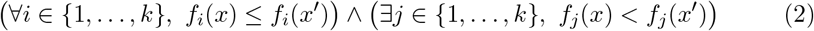

A solution *x* that is not dominated by any other solution *x*^*′*^ is called a Pareto optimal solution, and the set of all Pareto optimal solutions is called the Pareto front.

In this study, we aim to design novel proteins whose structures are similar to a known reference protein but exhibit low sequence similarity. This problem can be formulated as a multi-objective optimization problem with two objective functions: the sequence recovery rate *f*_recovery_(*x*), which measures the sequence similarity between a designed protein and the reference protein, and the structural similarity *f*_structure_(*x*), which measures similarity between a predicted structure of a designed protein and the known structure of the reference protein. This multi-objective optimization problem with *k* = 2 is expressed as Equation (3).

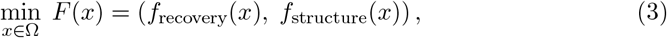

where Ω is the set of all amino acid sequences with the same residue length *n* as the reference protein. *f*_recovery_(*x*) is a function to measure the sequence similarity between the designed protein *x* and the reference protein *r*. It is defined as Equation (4).

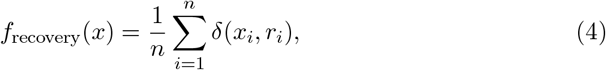

where *δ*(*x*_*i*_, *r*_*i*_) is an indicator function that returns 1 when the amino acids at position *i* match and 0 otherwise.

*f*_structure_(*x*) is a function representing the structural similarity between the predicted structure *AF* (*x*), the structure predicted by AlphaFold2 [15] from the amino acid sequence *x*, and the structure of the reference protein *R*. The AlphaFold2 predictions were performed via LocalColabFold [16]. As a metric for structural similarity, we use the Template Modeling score (TM-score) [17], which assigns higher values to more similar structures. *f*_structure_(*x*) is defined by inverting the sign of TM-score as in Equation (5) to treat this as a minimization problem.

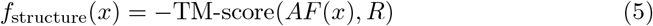

TM-scores were computed with USalign [18] using option 1, which superposes two structures by assuming that residues with the same index are equivalent between them, without performing structure-based alignment. Sequence-independent structural alignment can inflate similarity by matching discontinuous fragments, whereas our objective is global positional correspondence along the chain. Residue-by-residue alignment thus provides a more stable objective for optimization.

### 2.2. NSGA-II and mutation with ProteinMPNN

We used NSGA-II [14] to solve the multi-objective optimization problem. It is a widely used evolutionary algorithm that generates offspring from the parent population by genetic operations, such as crossover and mutation, and selects the next generation using non-dominated sorting and crowding distance to approximate the Pareto front. Instead of genetic operations in the original NSGA-II, we introduced a mutation operation that partially applies pMPNN to randomly selected three consecutive amino acids of a protein. By combining pMPNN and NSGA-II, we could incorporate the efficiency of generative AI algorithms into a conventional multi-objective optimization algorithm. The mutation operation using pMPNN is shown in Algorithm 1.

#### Algorithm 1 Mutation Operation Using ProteinMPNN

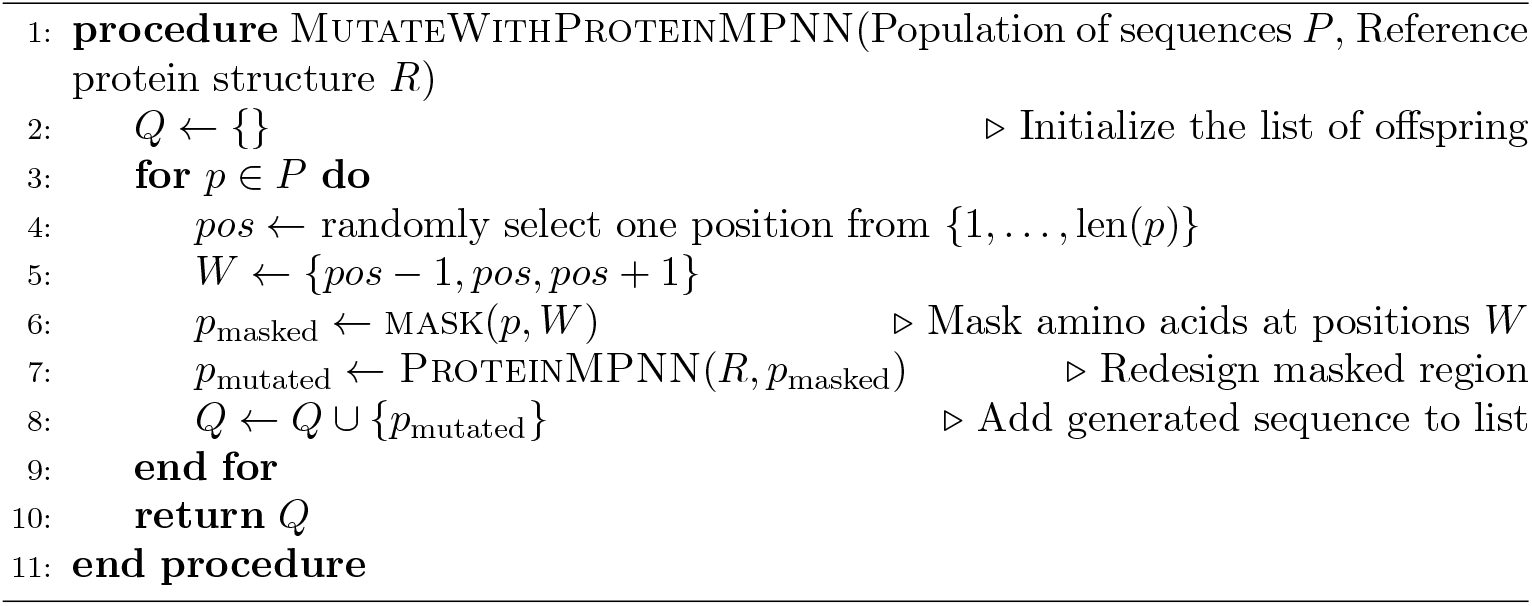

## 3. Results and Discussion

To evaluate the proposed algorithm, we conducted an experiment designing novel amino acid sequences using Top7 [19] as the reference protein. We executed the proposed method 24 times, each run starting from an initial population of amino acid sequences randomly generated with different seeds. Each sequence was represented by 92 characters, corresponding to the number of modeled residues in the Protein Data Bank entry of Top7 (PDB ID: 1QYS). The initial population size was set to 20, the number of generations was set to 500, and the temperature parameter of pMPNN was set to 0.3. As a baseline, sequence design using vanilla pMPNN was also performed on the same reference protein, with the temperature parameter set to the same value of 0.3. Note that the reference protein for TM-score calculation was the Top7 structure with selenomethionine residues replaced by methionine residues.

Figure 1 compares the sequence similarity to the reference protein between the proposed method and vanilla pMPNN. The proposed method produced proteins with lower sequence similarity to the reference, while maintaining comparable structural similarity. This result indicates that our method successfully designs proteins that are structurally similar but sequence-wise distinct from the reference.

**Figure 1.**
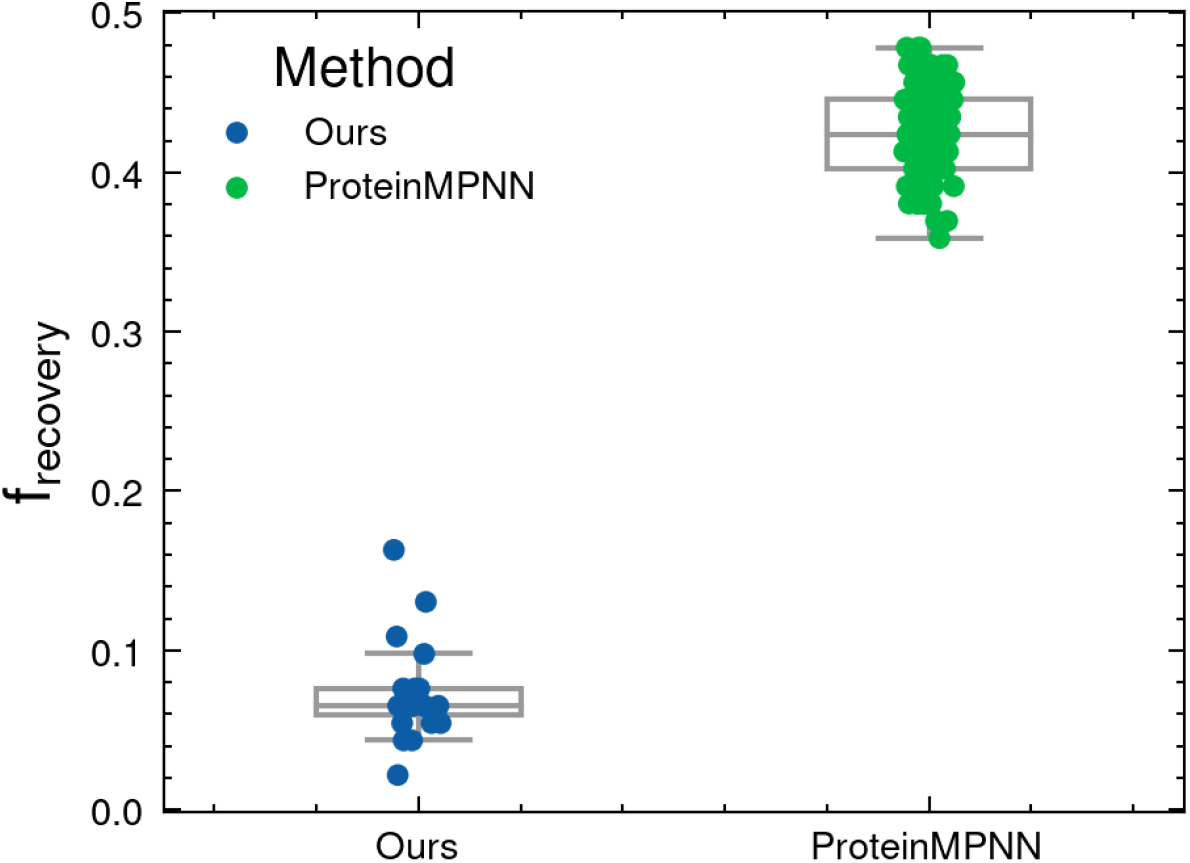
Comparison of sequence similarity for proteins obtained by the proposed method and by pMPNN. For the proposed method, the proteins with the lowest sequence similarity that satisfy TM-score ≥ 0.90 and pLDDT ≥ 90 are plotted for each run. The mean TM-score of the proteins designed by pMPNN was 0.95.

To evaluate the diversity of designed proteins, the protein sequence similarity network [20] is shown in Figure 2. A sequence similarity network is a network constructed based on the similarity between protein sequences and is used to visualize relationships and cluster structures within a set of sequences. In the proposed method, proteins derived from different seeds form multiple loose clusters and exhibit distributions that rarely overlap between seeds. In contrast, proteins generated by pMPNN mostly appear as a single dense cluster. This indicates that the proposed method can design more diverse proteins than pMPNN by changing the random seeds.

**Figure 2.**
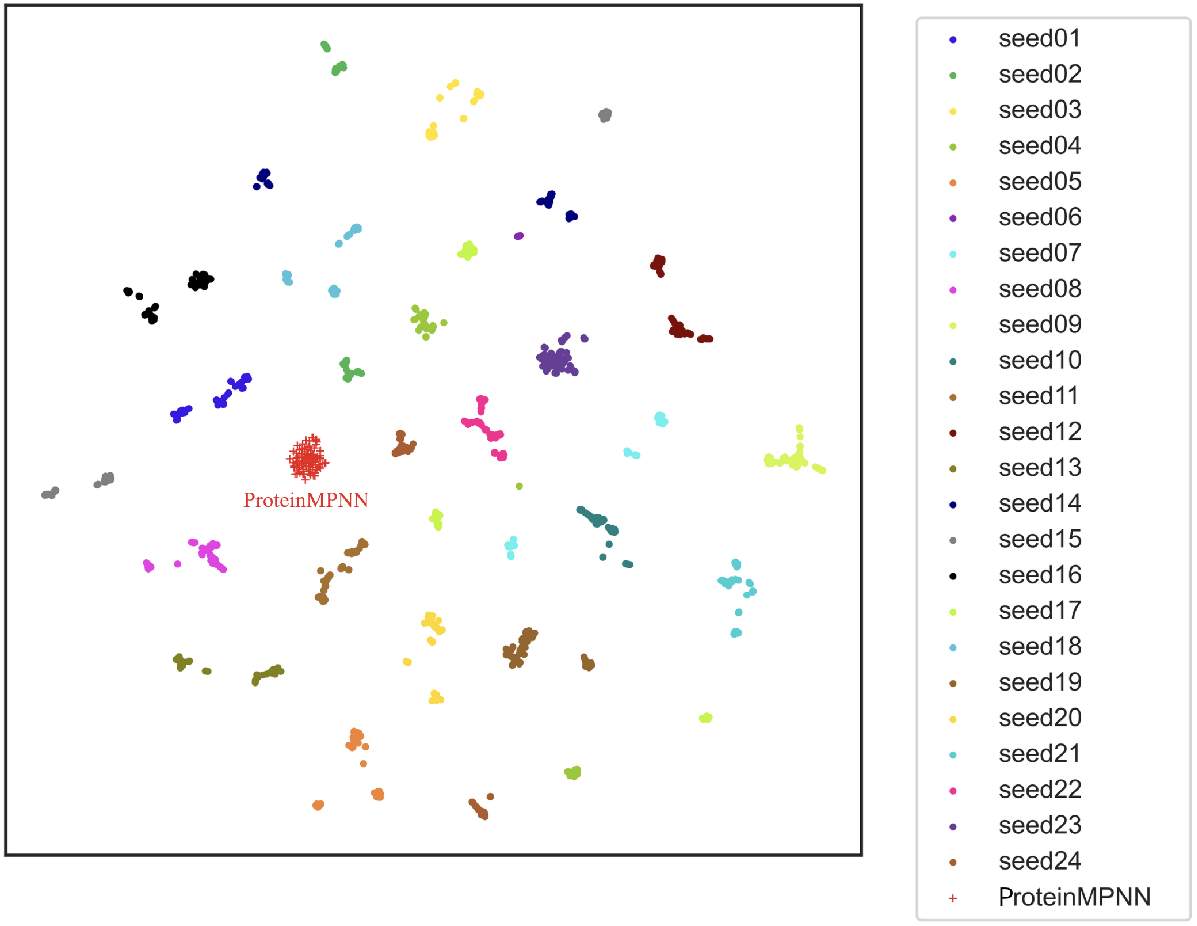
t-SNE plot of the Sequence Similarity Network (SSN) [20] of proteins generated by the proposed method and pMPNN. Points represent sequences generated by the proposed method from different random seeds that satisfy the criteria TM-score ≥ 0.9 and pLDDT ≥ 90. Up to 100 sequences are sampled from the Pareto front of each seed and are color-coded by seed. The central triangular cluster corresponds to the sequences generated by vanilla pMPNN.

To further analyze the effect of different random seeds, the Pareto fronts obtained by the proposed method using different random seeds are shown in Figure 3. While many seeds yielded high-quality protein designs with high structural similarity and low sequence similarity, some resulted in markedly inferior performance. This result highlights the importance of using multiple random seeds to achieve reliable performance.

**Figure 3.**
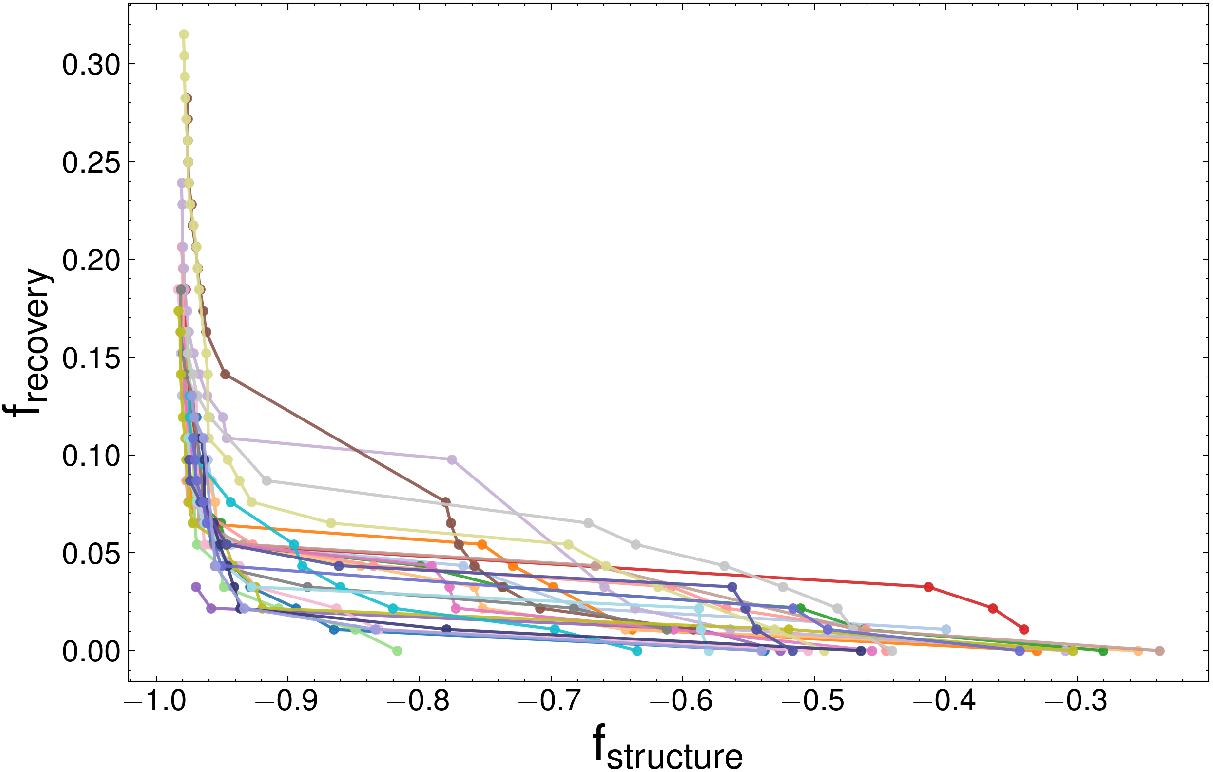
Pareto front obtained for each random seed.

Figure 4 shows the distribution of *f*_structure_ and *f*_recovery_ for proteins obtained from a single run of the proposed method starting from one random seed. In the initial generations, only proteins with TM-scores around 0.2 to 0.4 were obtained. However, as the generations progressed, proteins with higher TM-scores were obtained while maintaining low recovery values.

**Figure 4.**
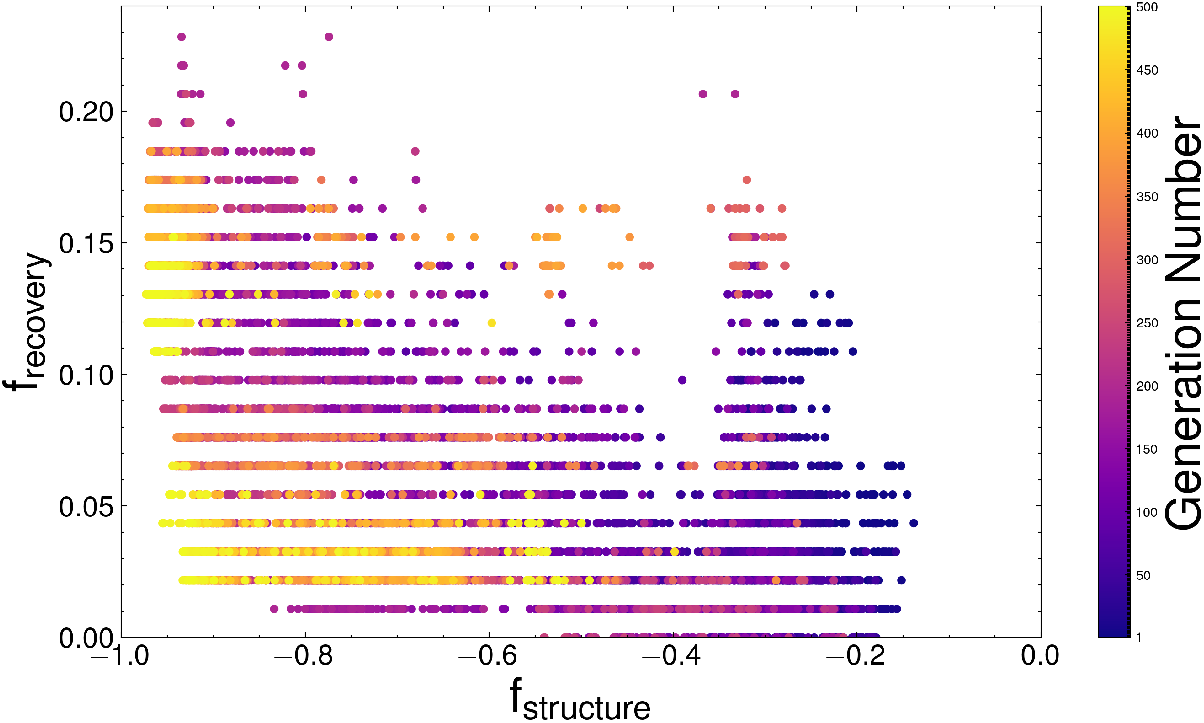
Distribution of *f*_structure_ and *f*_recovery_ for proteins obtained from a single run of the proposed method. Each protein is displayed in a different color according to the order in which it was obtained.

As a demonstrative example, we selected one protein designed by our method and superimposed its structures predicted by AlphaFold2, which was referenced during the design, and AlphaFold3 [21], which was not involved in the design process, onto the target structure using USalign (Figure 5). The structure predicted by AlphaFold2 showed a high pLDDT value of 93.12. Superimposition with the target structure (Figure 5a) yielded a TM-score of 0.96, indicating a very close match and supporting the effectiveness of the design. In addition, the AlphaFold3 prediction, despite being unused during design, had high confidence with a pTM score of 0.84 and a pLDDT value of 90.17. This structure also aligned closely with the reference, with a TM-score of 0.94 and RMSD of 0.93 °A upon superimposition (Figure 5b). This high structural agreement obtained from an independent prediction method suggests that the design is not simply overfitting to the AlphaFold2, but is likely capable of forming the intended target structure experimentally.

**Figure 5.**
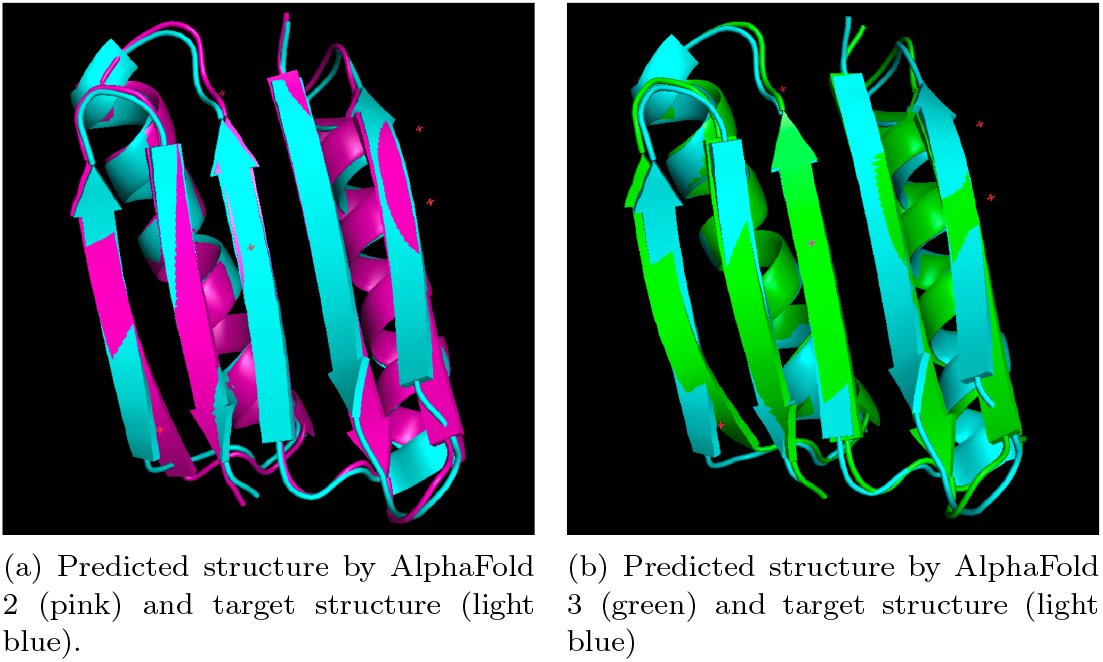
Predicted three-dimensional structure of the protein designed by the proposed method. PyMol [22] was used for visuallization.

## 4. Conclusion

In this paper, we proposed a method that combines pMPNN with a multi-objective optimization algorithm to design proteins with high structural similarity but low sequence similarity. Evaluation experiments showed that proteins designed by the proposed method exhibit greater sequence diversity compared to those designed by vanilla pMPNN. While this study used a similarity of predicted three-dimensional structure as the objective function, the framework of the multi-objective optimization method presented here can be applied to other objective functions as well. A major limitation of this method is the large number of evaluation function calls. When using objective functions that require experimental evaluation, it is necessary to further improve design efficiency by incorporating more efficient generative AI approaches, such as protein language models, to perform protein design with fewer evaluation calls. Furthermore, the proteins designed in this study have only been evaluated computationally. Experimental validation of the structural similarity of the designed proteins is part of our future work.

